# Democratizing Protein Language Models with Parameter-Efficient Fine-Tuning

**DOI:** 10.1101/2023.11.09.566187

**Authors:** Samuel Sledzieski, Meghana Kshirsagar, Minkyung Baek, Bonnie Berger, Rahul Dodhia, Juan Lavista Ferres

## Abstract

Proteomics has been revolutionized by large pre-trained protein language models, which learn unsupervised representations from large corpora of sequences. The parameters of these models are then fine-tuned in a supervised setting to tailor the model to a specific downstream task. However, as model size increases, the computational and memory footprint of fine-tuning becomes a barrier for many research groups. In the field of natural language processing, which has seen a similar explosion in the size of models, these challenges have been addressed by methods for parameter-efficient fine-tuning (PEFT). In this work, we newly bring parameter-efficient fine-tuning methods to proteomics. Using the parameter-efficient method LoRA, we train new models for two important proteomic tasks: predicting protein-protein interactions (PPI) and predicting the symmetry of homooligomers. We show that for homooligomer symmetry prediction, these approaches achieve performance competitive with traditional fine-tuning while requiring reduced memory and using three orders of magnitude fewer parameters. On the PPI prediction task, we surprisingly find that PEFT models actually outperform traditional fine-tuning while using two orders of magnitude fewer parameters. Here, we go even further to show that freezing the parameters of the language model and training only a classification head also outperforms fine-tuning, using five orders of magnitude fewer parameters, and that both of these models outperform state-of-the-art PPI prediction methods with substantially reduced compute. We also demonstrate that PEFT is robust to variations in training hyper-parameters, and elucidate where best practices for PEFT in proteomics differ from in natural language processing. Thus, we provide a blueprint to democratize the power of protein language model tuning to groups which have limited computational resources.

## 1 Introduction

The introduction of large pre-trained protein language models (PLMs) has transformed the computational modeling of protein sequence, structure, and function. These models are trained in an unsupervised manner on tens or hundreds of protein sequences, and they learn hidden representations which contain information about evolutionary constraints, chemical properties, secondary structure, and more [6]. These representations generalize broadly, which enables PLMs to be tuned to a wide variety of proteomic tasks. Typically, when a language model is tailored to a specific downstream task, the parameters of the pre-trained model are updated in a process known as fine-tuning (Figure 1a, top). However, as the size of foundation models increases, fine-tuning a model for a task of interest is increasingly computationally expensive, often with heavy GPU memory requirements. This puts such tuning out of the reach of many research groups, especially in academic, government, or startup environments. As increasingly large PLMs continue to be developed, such as the 6.4 billion parameter ProGen2 [22] or the 15 billion parameter ESM2 [20], the computational expense of fine-tuning will continue to be a pressing issue in proteomics.

**Fig. 1.**
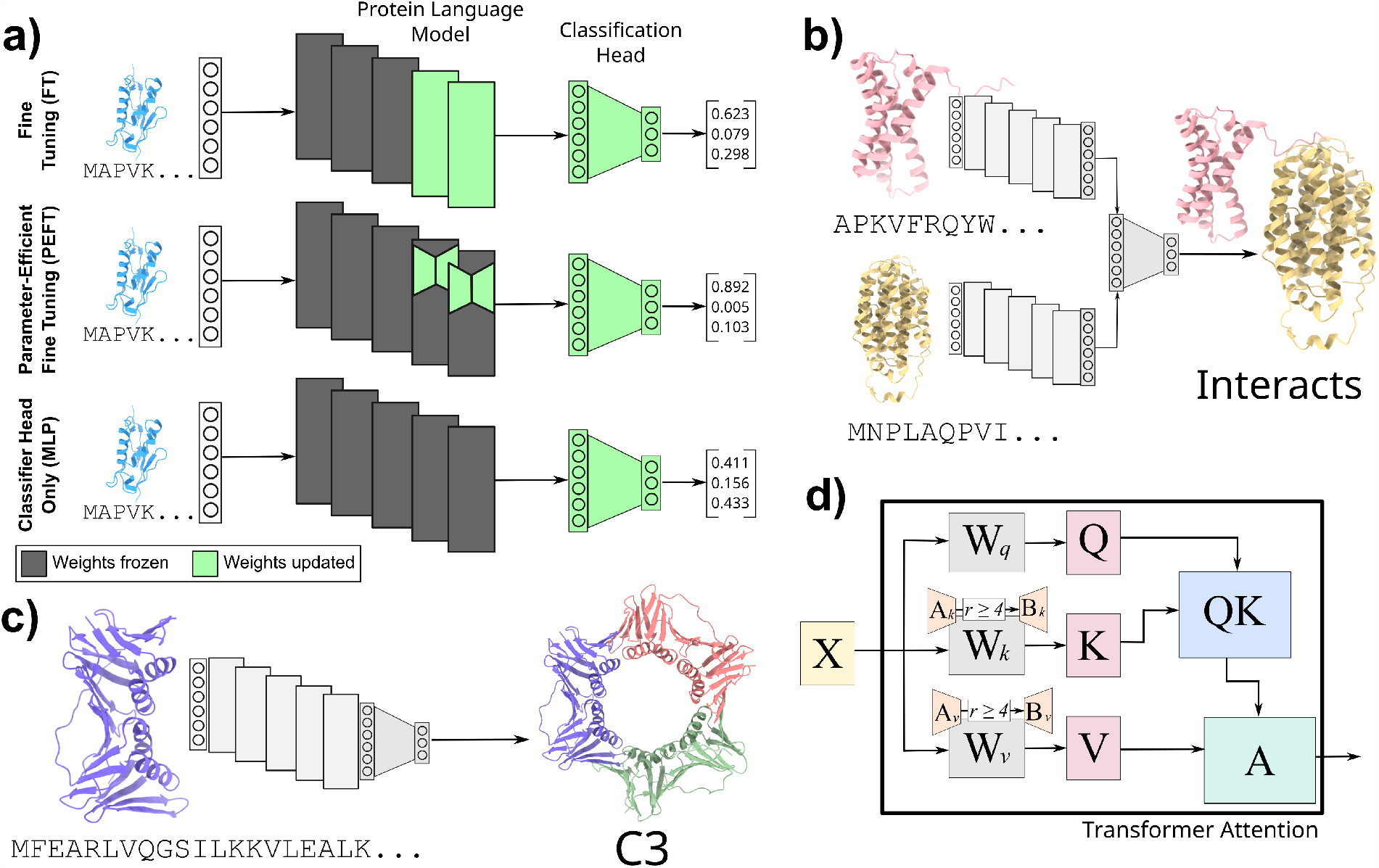
Bringing PEFT to proteomics. **(a)** The traditional paradigm for adapting protein language models to a specific downstream task is to fine-tune the parameters of the last *n* layers along with a new classification head (top, **FT**). Here, we introduce parameter-efficient fine-tuning (PEFT) to proteomics, tuning only the parameters of low-rank adapter matrices added to these final layers rather than the full weight matrices (middle, **PEFT**). We compare also with a baseline that uses the embeddings as-is and trains only the classification head (bottom, **MLP**) **(b)** We show that PEFT and MLP models achieve performance competitive with the state-of-the-art on predicting protein-protein interactions. **(c)** We also train PEFT models to predict the symmetry class of homooligomers, showing that the performance of PEFT models only slightly trails that of FT models yet uses several orders of magnitude fewer parameters. **(d)** We show the transformer self-attention with LoRA weight matrices added. We explore the hyper-parameter space of low-rank adapters, creating a blueprint for best applying LoRA to protein language models. Notably, this blueprint differs from that in NLP. We recommend adding adapters with rank at least 4 to the key and value weight matrices of the self-attention layers. *X*: input to attention head. *W*_*q,k,v*_: query, key, and value weights. *A*_*k,v*_, *B*_*k,v*_: newly added low-rank adapter weights. *Q, K, V* : query, key, and value representations. *QK*: intermediate product of *Q* and *K. A*: output of attention layer.

Natural language processing (NLP) has seen a similar increase in language model size, with the largest NLP models approaching or exceeding one trillion parameters. One alternative approach to fine-tuning is prompting, where the problem is presented in natural language and the model makes “zero shot” predictions [29], but it is not clear that such an approach translates directly to proteomics—how would one encode the question, “What is the structure of this protein?,” into a sequence of amino acids? Instead, we draw inspiration from methods developed for *parameter-efficient fine-tuning* (**PEFT**) in NLP. These methods add a small number (usually *<* 1% of total model size) of new parameters which are tuned, leaving the original model parameters untouched. Houlsby et al. [16] introduced adapters, which add parameters in serial to each transformer layer, allowing for every layer of the model to be trained using only a small number of parameters. The current state-of-the-art is low-rank adapters (LoRA), introduced by Hu et al. [17], which add two low-rank adapter matrices in parallel to the query and value weight matrices of the attention heads (Figure 1a, middle). These approaches rarely reach the performance of traditional fine-tuning (**FT**), but require significantly fewer resources.

Here, we introduce parameter-efficient fine-tuning methods to proteomics, demonstrating performance competitive with or exceeding traditional fine-tuning using parameter-efficient training on two important proteomic tasks— homooligomer symmetry prediction and protein-protein interaction prediction. Homooligomers (proteins that form complexes with copies of themselves) often adopt symmetric conformations. Predicting the symmetry that a homooligomer adopts is an important task in structural biology and protein design (Figure 1c, Section 1.1). We newly show that as in NLP, a PEFT model trails behind the performance of FT (*AUPR* = 0.400 vs. 0.489) but significantly outperforms baselines (*AUPR* = 0.238) and offers a much more compute-efficient alternative (Section 3.5).

We also train a PEFT model to predict protein-protein interactions (PPIs) from primary sequence, an important and well-studied problem (Figure 1b, Section 1.1). Surprisingly, we find that PEFT models outperform FT models (*AUPR* = 0.600 vs 0.577). Intrigued by this, we also tested the baseline model which trains an MLP classifier on embeddings from a frozen language model (Figure 1a, middle). We find that this method actually outperforms both tuning methods (*AUPR* = 0.684), demonstrating the continuing efficacy of simple models in proteomics. In fact, we show that both our PEFT and MLP models outperform the current state-of-the-art on a gold-standard benchmark from Bernett et al. [8] on several metrics (Section 3.2).

For the first time, we further create a blueprint for applying PEFT methods to protein language models on proteomics tasks by performing extensive experiments on LoRA hyper-parameter choices (Figure 1d). Contrary to what is recommended for NLP tasks, we find that adding LoRA adapters to only the key and value matrices of the transformer achieves optimal performance (Section 3.3), and that performance drops off as the rank of adapter matrices drops below 4 (Section 3.4). Our work shows that it is possible to achieve competitive performance with significantly fewer resources than traditional approaches, opening up the power of protein language model fine-tuning to academic labs, small biotech startups and other research groups lacking substantial computational resources.

### 1.1 Related Work

*Language Modeling in Biology*. While the first protein language models (Bepler & Berger, UniRep) used recurrent neural networks like the bi-LSTM [5,6,2], recent work has converged around masked language modeling and the transformer. Models like ProtBert, ProtT5, [13], and ESM [24] are transformers trained on massive sets of protein sequence data in an unsupervised manner. These models learn meaningful representations which can be applied to replace manual feature engineering, or computationally expensive evolutionary searches and construction of multiple sequence alignments. Most recently, ESM2 [20] represents the largest protein language model to date, with models as large as 15 billion parameters. While this is still shy of the largest natural language models, this represents a significant step up in the size of protein language models and their capacity for unsupervised representation learning. While language modeling has seen the most success in proteomics, this success has seen language models expand to other aspects of biology. Biochemistry language models learn representations of small molecules [25,15], most notably with ChemBERTa [11]. Likewise, the release of scGPT [12] has led to advancements in single-cell genomics, and language models have also seen direct clinical use, such as with the medical question answering model Med-PaLM [28].

#### Protein Structure

One of the longest standing challenges in computational biology is that of protein structure prediction. While proteins are modeled by language models as sequences of amino acids (primary structure), they fold into secondary (alpha helices, beta sheets) and more complex tertiary structures, which imbues them with a variety of functions. For nearly 30 years the Critical Assessment of protein Structure Prediction (CASP) has measured the ability to computationally predict the tertiary structure of a protein from its primary sequence. In 2020, AlphaFold2 [18], closely followed by RoseTTAFold in 2021 [4], presented a massive jump in performance, reaching near-experimental levels of accuracy. AlphaFold2 and RoseTTAFold use multiple sequence alignments (MSA) to incorporate evolutionary context into structure prediction, and recent methods like OmegaFold [34] and ESMFold [20] instead use pre-trained protein language models. While protein language model-based approaches have yet to reach the accuracy level of MSA-based approaches across the board, they nonetheless achieve extremely accurate performance and require only a single sequence. Because of this, they are much more computationally efficient, forgoing the need for the expensive MSA search step. In addition, these methods often outperform MSA-based methods on intrinsically disordered regions or proteins with little evolutionary context, and are competitive in terms of complex structure accuracy. Thus, protein language models represent an exciting step forward in tertiary structure modeling.

In addition to the structures that single chains fold into, proteins can form complexes known as quaternary structures comprising multiple protein chains. While methods like AlphaFold-Multimer [14] have attempted to fully model the structure of these complexes to moderate success, other methods have taken a different approach – focusing on the special case of homooligomeric proteins. In 2006, Levy et al. introduced 3DComplex [19], which categorizes protein complexes based on topology and symmetry. Recently, Schweke et al. introduced an atlas of protein homooligomerization [26], while QUEEN [3] attempts to predict the mutiplicity of such complexes.

#### Protein Interactions

Cellular function is driven by a complex interplay of interactions between proteins. Experimental approaches to discern those interactions require substantial wet lab resources and time, which motivates the need for computational approaches to model protein interaction. Models such as AlphaFold-Multimer [14] have recently been developed to predict the structure of interacting complexes [35]. Quaternary structure prediction is valuable if the pair is already known to interact, but often results in degenerate prediction for pairs which don’t interact, and due to the size of the model is difficult to scale to the whole-genome and all possible protein pairs. Methods like PIPR [10], D-SCRIPT [31], Topsy-Turvy [27] and RAPPPID [33] predict protein-protein interaction (PPI) solely from widely-available primary sequence and are fast enough to run at genome scale. Recent work has begun to close the gap between whole-genome interaction prediction and complex structure modeling [9,30], potentially unifying genome-scale PPI prediction with complex structure prediction. The Human Reference Interactome (HuRI) [21] remains the most complete experimentally-verified human protein interaction network.

## 2 Methods

### 2.1 Protein Language Model

We focused our efforts on ESM2, a transformer-based protein language model which is presently considered the state-of-the-art in protein language modeling [20]. ESM2 has several different model sizes, ranging from eight million to 15 billion parameters. For this study, we focused on the 650 million parameter version of ESM2 (Section S1.1).

For an amino acid sequence *X* = *x*_1_*x*_2_…*x*_n_, a PLM of dimension *d* returns a set of embeddings *E* ∈ ℝ^*d ×n*^ = *e*_1_*e*_2_…*e*_n_, *e*_i_ ∈ ℝ^*d*^. To standardize the size of representations for sequences of dynamic length, a pooling step needs to be undertaken. This is most commonly done either by averaging along the length of the sequences 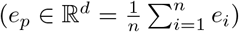 or by selecting the first token of the sequence, a non-amino acid token (`[clsf]`) created specifically for sequence classification. Here, we chose to take the former approach as it explicitly integrates signal across the length of the protein. We note that while this is a commonly used approach, how to best aggregate sequence-length representations into a fixed dimension embedding is an open problem in language modeling. Converting this pooled embedding into a binary (*Y* ∈ ℝ^2^) or multi-class (*Y* ∈ ℝ^18^) prediction requires an additional classification head. For the PPI prediction task, fixed-length embeddings were averaged before being passed to the classification head, as in Szymborski et al. [33]. For the symmetry prediction task, there is only a single protein, so embeddings were passed directly to the classification head. In this study, we tested two different prediction heads. The first, applied directly to the pre-trained *E* without fine-tuning, is a simple multi-layer perceptron (MLP), with the number and size of layers determined by grid search (Section 2.3, Section S1.3). The second is the `ESMClassificationHead` made available by the authors in the public HuggingFace repository, which consists of two dense layers with dropout and a tanh activation between the layers. We selected classification heads which were demonstrated to yield strong performance in previous work in order to minimize the need for hyper-parameter search in this space.

### 2.2 Parameter-Efficient Adaptation

Here, we chose to use LoRA [17], the most widely-adopted parameter-efficient fine-tuning method. LoRA adds two low-rank matrices *A* and *B* to each adapted weight matrix. Given weight matrix *W*∈ ℝ^*d×k*^, LoRA adds new parameters *A* ∈ ℝ^*r×k*^, *B* ∈ ℝ^*d×r*^, *r << d, k*. The normal forward pass of the layer given input *x* ∈ ℝ^*k*^ is *h* = *Wx*, and the forward pass with the LoRA adaptation is *h* = *Wx* + *BAx*. Only the weights of *A, B* are updated during back-propagation, while the weights of *W* are frozen. *BAx* is scaled by the quantity 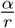, where *α* is a hyper-parameter which is held constant in the original report. While *A* is initialized with a random Gaussian distribution, by initializing *B* = 0 the first forward pass of the model is equivalent to the pre-trained model without adaptation. Following the recommendations of the original paper, we initially apply LoRA only to the query and value matrices of the attention head. We explore different combinations of weight matrix adaptation in Section 3.3 and different values of the rank *r* in Section 3.4.

### 2.3 Training and Implementation

All PEFT and FT models were implemented in PyTorch, using the HuggingFace implementations of ESM2 from the `transformers` package and LoRA from the `peft` package. Models were trained on NVIDIA V100 GPUs with 32GB of memory. We used the binary cross-entropy with logits loss to compute error, with an L2 weight decay of 0.01. Model weights were optimized via back-propagation using the Adam optimizer and a cosine decay with restarts learning rate schedule (initial learning rate 0.001). Models were trained with an epoch size of 16, 384, with an on-device batch size of 4 and gradient accumulation every 16 steps, for an effective batch size of 64. PEFT and FT models were trained for 40 epochs and the best model based on validation AUPR was chosen for testing. Except for where otherwise specified, we used LoRA *r* = 8, LoRA *α* = 32, LoRA dropout *p* = 0.1.

The hyper-parameters for the MLP on language model embeddings with frozen weights were chosen by a grid search (implemented in scikit-learn, Section S5). The best performing model had two hidden layers with sizes (64, 64) and ReLU activations, and were optimized with the Adam optimizer for 2000 iterations with a tolerance of 0.0001 and an adaptive learning rate initialized at 0.01.

### 2.4 Benchmark Data – PPI

While creating train/test splits based on filtering homologous proteins is common in machine learning for proteomics, the binary nature of PPI prediction presents a unique challenge because data leakage can still occur if only one protein of an interacting pair appears in both sets. If a so called “hub” protein with many interactions appears in both the training and test set, models can learn that this specific protein is likely to have positive interactions. Then, test set performance will be inflated even if nothing is learned about the actual pairwise interactions. After noting pervasive biases in previous benchmarks relating to sequence similarity and node degree, Bernett et al. [8] introduced a new gold standard data set for benchmarking PPI. The splits introduced in this benchmark apply a more stringent notion of sequence similarity for pairwise problems as introduced by Park and Marcotte [23], splitting by C3 similarity. In addition, both the positive and negative data sets are balanced with regard to node degree; as a consequence models cannot learn that proteins *in general* interact just because they are high degree. This data set consists of 163,192/59,246/52,035 training/validation/test edges, with an 1:1 ratio of positives to negatives.

### 2.5 Benchmark Data – Homooligomer Symmetry

The multiplicity and symmetry prediction task is formulated as a multi-class prediction problem. Given a protein chain, we classify it as one of 17 symmetry classes *C*1, *C*2, *C*3, *C*4, *C*5, *C*6, *C*7 − *C*9, *C*10 − *C*17, *D*2, *D*3, *D*4, *D*5, *D*6 − *D*12, *H, O, T, I* (or “Unknown”). The *C* classes correspond to cyclic symmetries, the *D* classes to dihedral symmetries, and H, O, T, and I to helical, octahedral, tetrahedral, and icosahedral symmetries respectively. More extensive detail on different symmetry classes can be found within 3DComplex [19]. Protein sequences and structures were obtained from the Protein Data Bank (PDB) [7], as were their labels. Sequences were clustered at 30% sequence similarity and 80% coverage using MMseqs2 [32], and these clusters were used to define train, validation, and test splits. This method of splitting ensures that no two sequences that are highly similar will be in both a training and evalution set, which would allow the model to memorize sequence similarity rather than learning properties of sequence and structure that correspond with symmetry. This data consists of 370,986/46,833/102,978 training/validation/test edges. The support for each class in the text set is described in Table S2.

We tracked performance of models through multiple metrics. It is difficult to pin performance to a single number with a multi-class classification, especially when the support for different classes is highly imbalanced. We compute the accuracy, F1 score, MCC, average precision (AUPR), precision, recall, and specifity for each class, as well as metrics averaged over each class. Note that because class support is highly variable (Section S2.1) and resulting model performance varies widely, we use an unweighted (“macro”) average to capture performance broadly across all classes.

## 3 Results

### 3.1 Reduced memory usage of PEFT enables deeper fine-tuning

The most common approach to fine-tuning is to unfreeze the weights of the last *n* transformer layers, which we compare with adding LoRA adapter weights to the last *n* layers. Table 1 shows the maximum GPU memory used by parameter-efficient fine-tuning (PEFT) or traditional fine-tuning (FT) for the last *n* layers of an ESM2 model trained on PPI prediction or homooligomer symmetry prediction. Both PEFT and FT eventually overflow available GPU memory as the number of layers tuned increases; tuning all layers of the model requires additional model parallelism with either approach. However, this parameter-efficient approach allows for training of deeper layers of the model while staying within a lower memory budget. Because the PPI task requires storing the compute graph for a pair of proteins, the marginal impact of PEFT methods on memory consumption is less than for the symmetry task, where using LoRA to adapt weight matrices allows for several additional layers to be trained. All experiments were performed on a GPU with 32GB of memory, with a batch size of 4 and maximum sequence length of 1024.

**Table 1.**
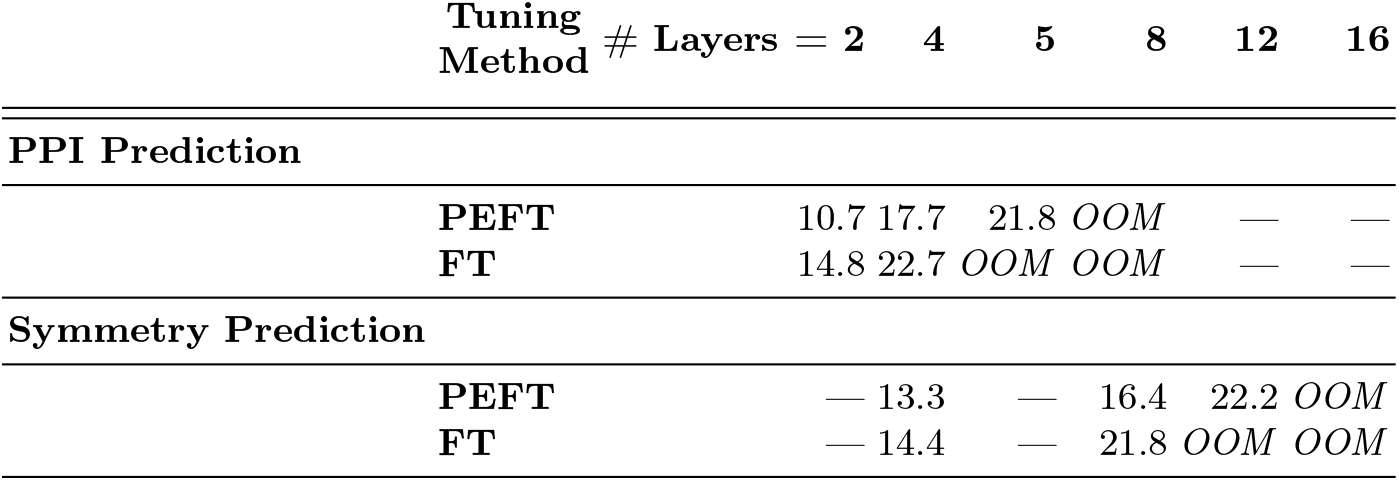
PEFT training requires reduced GPU memory. We compare the maximum GPU memory usage of parameter-efficient fine-tuning vs. traditional fine-tuning different numbers of transformer layers. All values are reported in GB. *OOM* indicates that the run was killed due to running out of GPU memory. Parameter-efficient fine-tuning enables adaptation of deeper model layers within the same sized GPU. More layers are able to be adapted for the symmetry prediction task since it requires only a single protein, rather than a pair.

| Tuning Method | # Layers = | 2 | 4 | 5 | 8 | 12 | 16 |
| --- | --- | --- | --- | --- | --- | --- | --- |
| **PPI Prediction** |  |  |  |  |  |  |  |
| **PEFT** |  | 10.7 | 17.7 | 21.8 | *OOM* | — | — |
| **FT** |  | 14.8 | 22.7 | *OOM* | *OOM* | — | — |
| **Symmetry Prediction** |  |  |  |  |  |  |  |
| **PEFT** |  | — | 13.3 | — | 16.4 | 22.2 | *OOM* |
| **FT** |  | — | 14.4 | — | 21.8 | *OOM* | *OOM* |

### 3.2 Efficient classifiers achieve state-of-the-art PPI prediction

We use parameter-efficient fine-tuning to train an ESM2 model (PEFT) to predict protein-protein interactions from sequence on the benchmark data set from Bernett et al. [8]. We compare with a model that is fine-tuned in the traditional way (FT), and with a baseline that trains a classifier on sequence embeddings from the ESM2 model with frozen weights (MLP). See Figure 1a for an overview of these three approaches. In addition, we compare to the best prior scores from Bernett et al. [8]—either Topsy-Turvy [27], or SVM-PCA, a baseline constructed by Bernett et al. [8] which trains a support vector machine on PCA-reduced sequence similarity vectors. Note that for each benchmark metric, we selected the best score across *all* methods evaluated, and that no single method achieved the “Best Prior” performance across the board, so this is a significantly higher threshold than comparing to any single method.

Table 2 shows the performance of these models. Both the PEFT and MLP models achieve stronger performance than the FT model, despite having several orders of magnitude fewer parameters. In fact, the MLP model achieves the best overall accuracy and AUPR on these benchmarks. Both the MLP and PEFT models actually *outperform* the “Best Prior” methods in these metrics, as well as F1 and MCC. Surprisingly, both the MLP and PEFT models outperform the “Best Prior” methods in these metrics, as well as F1 and MCC. These results suggest that larger, more complex models are not always better, at least for the PPI prediction task (see also Section S1.1). Both models use less GPU memory than the FT model (MLP substantially so) and the MLP model requires significantly less training time.

**Table 2.**
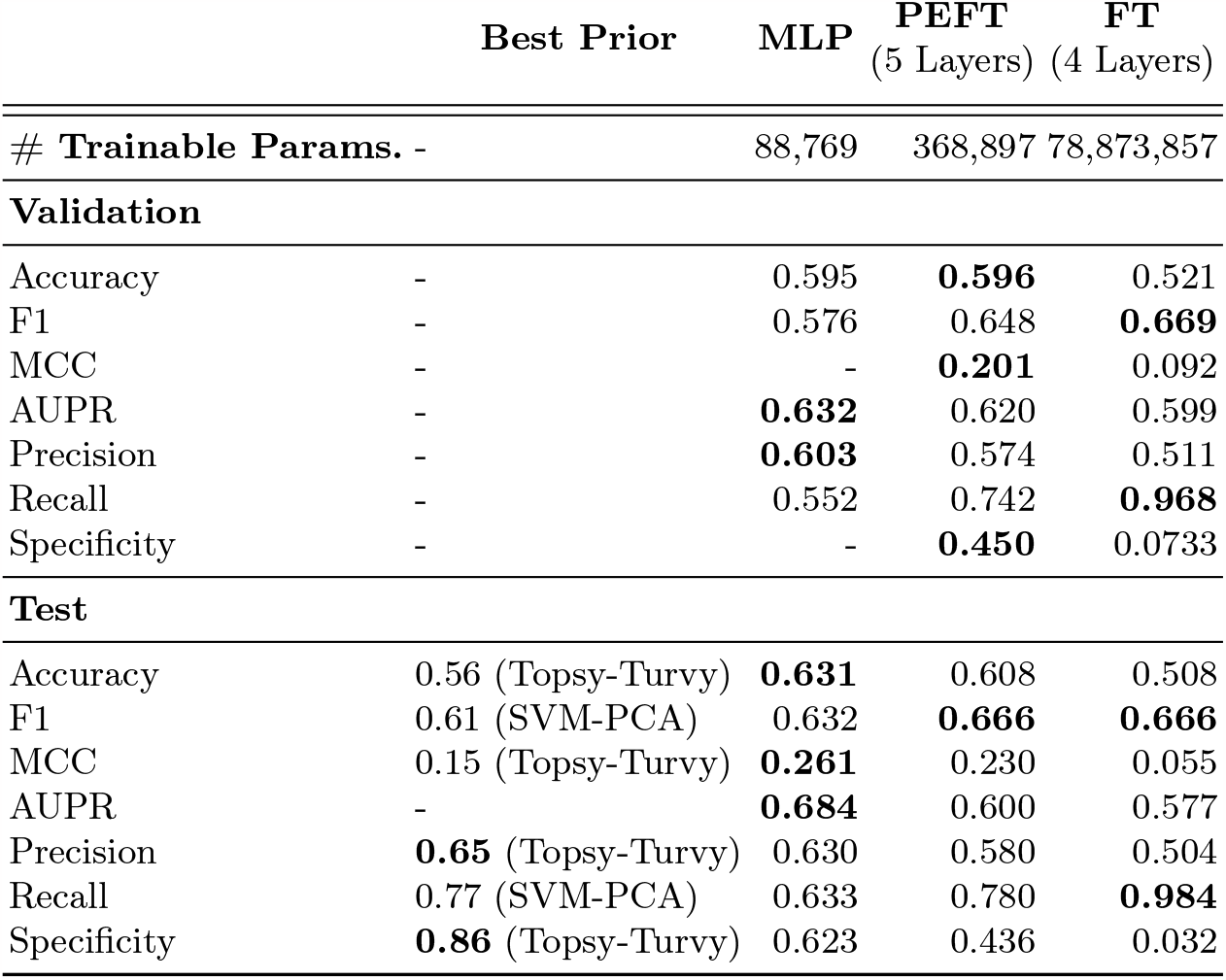
Applying PEFT to train models for protein-protein interaction. We trained multiple variants of ESM2 to predict protein-protein interactions, and evaluate using the benchmark data sets from Bernett et al. [8]. **MLP** indicates a multi-layer perceptron trained on embeddings from a frozen model, while **PEFT** and **FT** indicate parameter-efficient fine-tuning and traditional fine-tuning of the transformer layers. Due to the reduced memory footprint of PEFT, we were able to fine-tune an additional layer. Simply using ESM2 embeddings with an MLP classifier outperforms the best prior methods across most metrics. The PEFT model achieves increased recall and F1 score compared to the MLP model, and also performs the best prior. Permitting all parameters to be fine-tuned yields not only worse performance, but training is much less stable; over the course of training, validation performance rapidly fluctuates between extremely high recall/near zero specificity and the reverse (Supplementary Figure S2). By contrast, validation performance with PEFT training is significantly more stable.

We note that FT model has an extremely high recall, but a near-zero specificity. As we show in Supplementary Figures S1, S2, despite relatively stable behavior of the training and validation loss, validation recall and specificity of the FT model is highly variable—fluctuating wildly between zero and one over the course of training. In contrast, PEFT training is much more stable. This suggests that the restriction of the low rank updates limits the variation in model weights and predictions, decreasing the importance of selecting the *exact* epoch to stop training to achieve the desired performance.

### 3.3 Which matrices in the transformer layers should be adapted for protein modeling**?**

In their original manuscript on LoRA, Hu et al. [17] show that for natural language models, adding low-rank adapter matrices to only the query and value weights (*W*_Q_, *W*_V_) of the attention heads yields the best tradeoff of performance and parameter-efficiency. However, the space of natural language is not necessarily the same as that of protein sequence, and we sought to evaluate to what extent the choice in adapted weight matrices affects performance. Table 3 shows that while performance is relatively robust, adapting the key (*W*_K_) and value (*W*_V_) matrices results in the best overall performance. We note that the value matrix achieves similar results while also being a more parameter-efficient. Thus, that could be the best choice for parameter-efficient fine-tuning of PLMs, if memory constraints are especially tight. When applying LoRA to protein language models, we recommend adding adapter matrices to the key and value weight matrices.

**Table 3.** Adaptation of different sets of transformer weights. Hu et al. [17] recommend adding low-rank adapters to the query and value matrices of the attention heads for natural language. We investigate whether the same recommendation holds for the space of protein language, specifically on the task on protein-protein interaction prediction. We report the validation AUPR, which was used to select the best epoch for test set evaluation, and the AUPR, accuracy, F1 score, MCC, precision, recall, and specificity on the test set for all different combinations of the query (*W*_Q_), key (*W*_K_), and value (*W*_V_) matrices. We find that adapting the *W*_K_ and *W*_V_ together yields the best performance. Adapting only the value matrix also performs well and uses fewer parameters.

| Weight Matrix | Val.<br>AUPR | AUPR | Acc. | F1 | MCC | Prec. | Rec. | Spec. |
| --- | --- | --- | --- | --- | --- | --- | --- | --- |
| $W_Q$ | 0.536 | 0.529 | 0.516 | 0.430 | 0.034 | 0.524 | 0.364 | 0.669 |
| $W_K$ | 0.565 | 0.562 | 0.544 | 0.433 | 0.095 | 0.572 | 0.348 | **0.739** |
| $W_V$ | 0.612 | 0.610 | **0.605** | **0.650** | **0.216** | 0.583 | **0.735** | 0.474 |
| $W_Q, W_K$ | 0.590 | 0.576 | 0.564 | 0.582 | 0.129 | 0.559 | 0.607 | 0.521 |
| $W_Q, W_V$ | 0.619 | 0.617 | 0.599 | 0.633 | 0.201 | 0.583 | 0.692 | 0.506 |
| $W_K, W_V$ | **0.637** | **0.633** | 0.601 | 0.630 | 0.205 | **0.587** | 0.680 | 0.522 |
| $W_Q, W_K, W_V$ | 0.628 | 0.613 | 0.603 | 0.639 | 0.210 | 0.585 | 0.704 | 0.502 |

### 3.4 Impact of LoRA rank on PPI prediction

The rank *r* of the newly added LoRA weight matrices plays an important role in the performance of PEFT models and the memory that they require. Hu et al. [17] show that LoRA remains effective at extremely low ranks, with competitive performance even when *r* = 1. However, the effectiveness of different rank values is dependent on both the intrinsic dimension of the language and the task [1], and robustness in rank variance will not necessarily hold for proteomic sequences and inference tasks. In Table 4, we show the results of training PEFT models with LoRA rank *r* = 1, 2, 4, 8, 64 (following Hu et al. [17]) to predict protein-protein interactions. We find that the best model performance is achieved at *r* = 4, but that higher ranks yield similar performance. For ranks less than 4, model performance is still strong, but is noticeably reduced. We perform the same experiments on homooligomer symmetry prediction, with similar findings—although *r* = 8 is better for this task (Table S3).

**Table 4.** Variation in rank is tolerated, but performance degrades at too low a rank. Hu et al. [17] show that LoRA is robust to rank *r* values as low as one. We test whether this robustness holds for protein language and on the PPI prediction task. Test set performance remains relatively strong across all ranks tested and actually peaks at *r* = 4, but there is a noticeable drop-off in performance for *r* = 1, *r* = 2. We recommend using a rank of at least 4 when fine-tuning protein language models.

| Rank | Val.<br>AUPR | AUPR | Acc. | F1 | MCC | Prec. | Rec. | Spec. |
| --- | --- | --- | --- | --- | --- | --- | --- | --- |
| 1 | 0.621 | 0.600 | 0.592 | 0.635 | 0.190 | 0.575 | 0.710 | 0.474 |
| 2 | 0.619 | 0.586 | 0.595 | 0.614 | 0.190 | 0.586 | 0.644 | **0.545** |
| 4 | **0.644** | **0.633** | **0.604** | **0.658** | **0.219** | 0.579 | 0.761 | 0.447 |
| 8 | 0.637 | **0.633** | 0.601 | 0.630 | 0.205 | 0.587 | **0.680** | 0.522 |
| 64 | 0.638 | 0.623 | 0.601 | 0.630 | 0.205 | **0.588** | 0.679 | 0.523 |

### 3.5 Predicting homooligomer symmetry with PEFT

We additionally train PEFT, FT, and MLP models to predict homooligomer symmetry. While this task also involves learning representations that capture protein structure, it is fundamentally different from the PPI task because it requires learning on only a single protein, rather than a pair. Thus, we capture the performance of PEFT training on diverse proteomic tasks. Contrary to what we find for PPI prediction, the relationship between the MLP, PEFT, and FT models is much closer to what would be expected based on model complexity. As we show in Table 5, both the PEFT and FT models significantly outperform the MLP model on all metrics. Traditional fine-tuning still yields the best performance, but parameter-efficient fine-tuning using LoRA is a viable alternative—performance is within 10-15% of the FT model, and the model uses 3 orders of magnitude fewer parameters. The PEFT model actually achieves the best MCC and specificity of the three.

**Table 5.** Applying PEFT to train models for homooligomer symmetry. We trained multiple variants of ESM2 to predict the symmetry of homooligomers from primary sequence. **MLP** indicates a multi-layer perceptron trained on embeddings from a frozen model, while **PEFT** and **FT** indicate parameter-efficient fine-tuning and traditional fine-tuning of the transformer layers. Due to the reduced memory footprint of PEFT, we were able to fine-tune four additional layers (12 vs. 8). While traditional fine-tuning still yields the best performance, PEFT is competitive across many metrics and significantly outperforms training only the classification head.

|  | **MLP** | **PEFT**<br>(12 Layers) | **FT**<br>(8 Layers) |
| --- | --- | --- | --- |
| # Trainable Params. | 89,857 | 657,810 | 157,585,810 |
| Accuracy | 0.206 | 0.393 | **0.413** |
| F1 | 0.228 | 0.405 | **0.455** |
| MCC | 0.299 | **0.464** | 0.460 |
| AUPR | 0.238 | 0.400 | **0.489** |
| Precision | 0.341 | 0.477 | **0.618** |
| Recall | 0.206 | 0.393 | **0.413** |
| Specificity | 0.960 | **0.970** | **0.970** |

Unlike the binary prediction task of PPI prediction, homooligomer symmetry prediction is a multi-class problem, where we consider 17 different possible symmetries (and an 18th “Unknown” class). This makes it more difficult to summarize model performance into a single number—especially because the number of examples of different classes in the test set is so variable (Supplementary Table S2). The numbers reported in Table 5 are macro averages across all classes, but in Figure 2 we show the accuracy, AUPR, and F1 score of the three models broken down by true class. These results make it clear that the improved performance of the PEFT and FT models comes primarily from their increased ability to learn about rare classes. On common classes like C1, C5, D2, and I, the MLP model achieves accuracy competitive with the PEFT model and often even the fine-tuned model. However, the larger complexity of the FT model allows it to also learn about classes like C3, C4, D6-D12, and O significantly better than the less complex models.

**Fig. 2.**
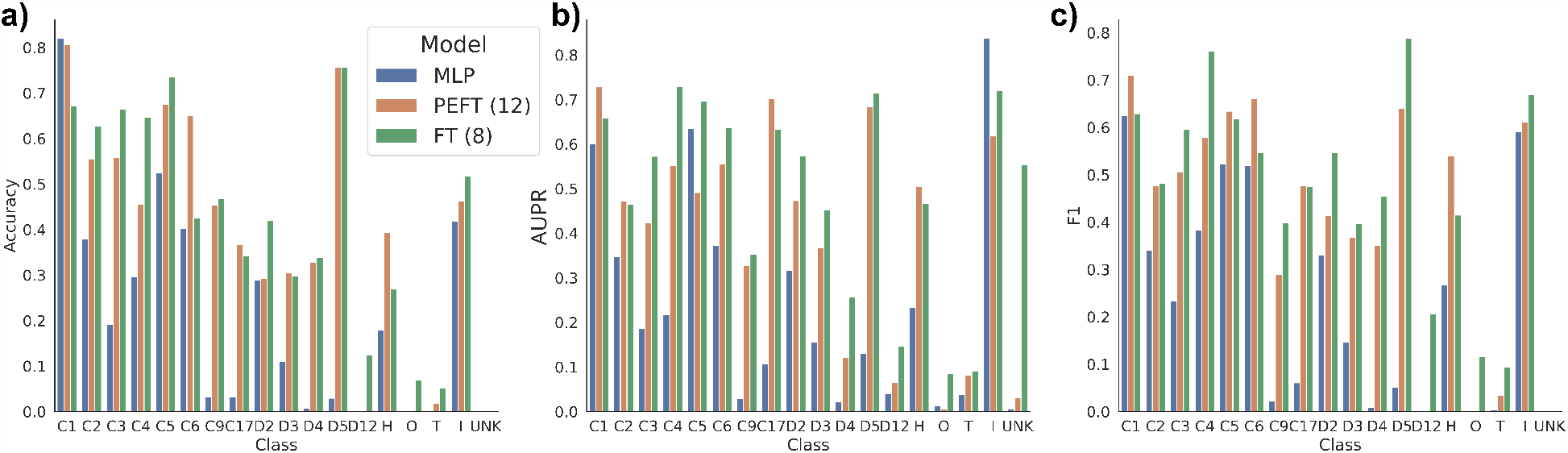
Breakdown of model performance by homooligomer symmetry class. Looking at only a single set of metrics can be helpful to get an aggregate idea of model performance, but hides complexity in the case of a multi-class classification task such as homooligomer symmetry prediction. Here, we show the per-class accuracy **(a)**, AUPR **(b)**, and F1 **(c)** of a classifier trained with frozen ESM2 embeddings (**MLP**), a model fine-tuned with LoRA (**PEFT**, 12 layers), and a model where all paramters in the final 8 layers were fine-tuned (**FT**). **MLP** performance is unsurprisingly relatively strong for high-support “easy” classes (e.g. C1, C5, D2, I), but substantially worse for rarer classes (e.g. C7-9, C10-C17, D4, D5). The **PEFT** model is competitive for most classes, but generally lags behind the performance of the **FT** model.

### 3.6 Visualizing attentions after fine-tuning

In Figure 3, we show the impact of parameter-efficient fine-tuning on attention (*A* matrix from Figure 1d). Here, we look at the PEFT model trained for PPI prediction from Table 2. We visualize attention from the five LoRA-adapted layers averaged across all heads. Figure 3a shows attention from the pre-trained model for NADH dehydrogenase 1 *β* subcomplex subunit 1 (UniProt ID: O75438) (Figure 3c). Attention is concentrated along the diagonal. In contrast, after PEFT with LoRA weights the attention is much more diffuse across the length of the protein, especially in the later transformer layers (Figure 3b). This suggests that for PPI prediction, the LoRA weights allow for more distant amino acids to attend to one another. We show one representative example here, but this diffusion of attention occurs broadly; we show another representative example (NADH dehydrogenase 1 *β* subcomplex subunit 10, UniProt ID: O96000, which interacts with O075438 in our test set) in Figure S6. For symmetry, this effect is less pronounced, perhaps because it is only a single-chain task (Figure S7).

**Fig. 3.**
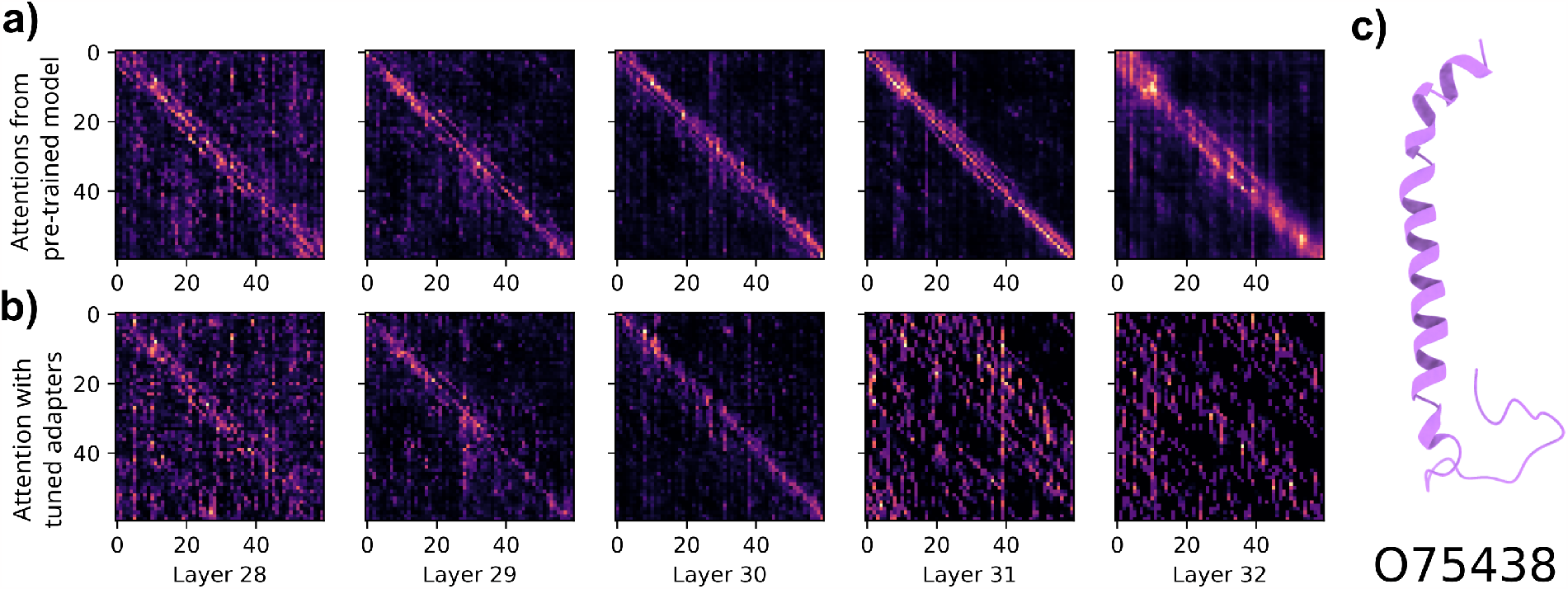
Visualizing attention matrices. **(a)** Attentions for NADH dehydrogenase 1 *β* subcomplex subunit 1 (UniProt ID: O75438) using the pre-trained ESM2. **(b)** Attentions for the same protein after parameter-efficient fine-tuning. **(c)** Structure of O75438. We find that PEFT weights result in attention that is more spread out across the length of the protein when trained for PPI prediction.

## 4 Discussion

In this manuscript, we bring parameter-efficient fine-tuning methods to protein language models. We show that PEFT models achieve comparable or better performance than traditional fine-tuning with a reduced memory footprint. This reduced memory allows for tuning of deep transformer layers, resulting in not only better performance but more stable training. Our work democratizes protein language models by charting a path for fine-tuning these models using substantially fewer parameters and less GPU memory. As the scale of PLMs continue to increase, it will become increasingly important to use these and other approaches when traditional fine-tuning is computationally infeasible.

Additionally, while approaches from natural language processing have so far transferred quite well, the distributions and intrinsic dimensions of protein language are clearly different than natural language. We show one consequence of this; our recommendation for LoRA rank and which attention matrices to adapt differs from the original conclusions of Hu et al. [17]. It may be necessary in the future to develop parameter-efficient approaches specific to protein language. There are still several open questions in this area—for example, while LoRA is the most performant PEFT method for natural language, other PEFT methods exist which may better suit the space of protein language. In addition, the difference in PEFT performance between PPI and homooligomer symmetry prediction tasks suggests that the efficacy of PEFT methods may be more variable, and further exploration on more diverse proteomic tasks would help to answer this question.

This work also shows the limitations of scale in protein language modeling. Not only are PEFT and MLP models able to achieve state-of-the-art performance on PPI prediction, but we show that embeddings from the 650 million parameter version of ESM2 outperform those from the 3 billion parameter version. While larger models certainly enable better performance, they do not guarantee it; we emphasize the continued need for models which can be easily trained and run by even small research groups. Especially with the cost and environmental impact of large-scale language model training and tuning, parameter-efficient tuning or methods which use frozen embeddings should remain a viable alternative for tailoring foundation models to a specific task, and should be tested prior to using traditional fine-tuning approaches.

## Acknowledgements

Thanks to Zhongqi Miao and Zalan Fabian for helpful discussions regarding parameter-efficient fine-tuning for large language models. Thanks to Bowen Jing, Lena Erlach, and Shuvom Sadhuka for feedback on the manuscript.

