## Supplementary Material for "Democratizing Protein Language Models with Parameter-Efficient Fine-Tuning"

### S1 Protein-Protein Interaction Prediction

#### S1.1 650M vs. 3B ESM2 Model

Typically in language modeling, larger models yield better performance leading to training or increasingly large models. We trained a multi-layer perceptron classifier (MLP) on embeddings from both 650M and 3B parameter models with frozen weights to predict protein-protein interactions. We found that despite having 4.5x fewer parameters, the 650M parameter model actually performs slightly better (Table S1). This indicates that even with reduced compute capacity available, smaller foundation models may be sufficient to achieve good performance on proteomics tasks—and that the limiting factor for performance may not simply be the scale of models. Consequently, all results presented elsewhere in this manuscript use the 650M parameter version of ESM2. We discuss the question of foundation model size in Section 4.

**Table S1.** Test set performance of a MLP classifier trained on pooled embeddings from the 650 million and 3 billion parameter versions of ESM2 with frozen weights. While the two models are competitive, the 650M parameter version outperforms the larger 3B parameter version in accuracy, MCC, AUPR, precision, and specificity. The 3B parameter model achieves a higher F1 score and recall.

| | Accuracy | F1 | MCC | AUPR | Prec. | Rec. | Spec. |
|---|---|---|---|---|---|---|---|
| **650M** | **0.631** | 0.632 | **0.261** | **0.684** | **0.630** | 0.633 | **0.623** |
| **3B** | 0.607 | **0.650** | 0.221 | 0.656 | 0.586 | **0.730** | 0.484 |

#### S1.2 PEFT and FT Training and Validation Curves

We show training and validation loss curves, as well as validation AUPR curves, over training in Figure S1. We show validation recall and specificity curves in Figure S2. In Figure S3, we show training and validation loss curves, as well as validation AUPR curves, for all different combinations of Q/K/V matrices tested in Table 3. In Figure S4, we show training and validation loss curves, as well as validation AUPR curves, for LoRA rank $1, 2, 4, 8, 64$ as tested in Table 4.

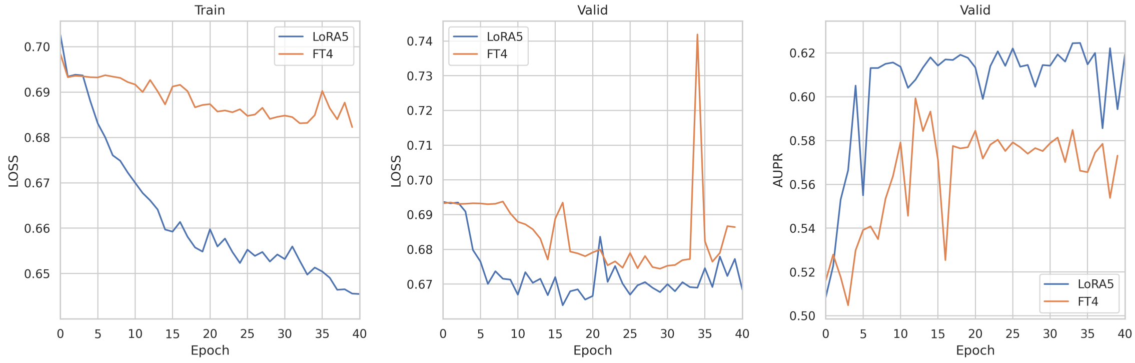


**Fig. S1.** Training and validation loss curves, AUPR curves for FT (4 layers) and PEFT (5 layers) from Table 2. Note that all other training parameters were held constant between these runs.

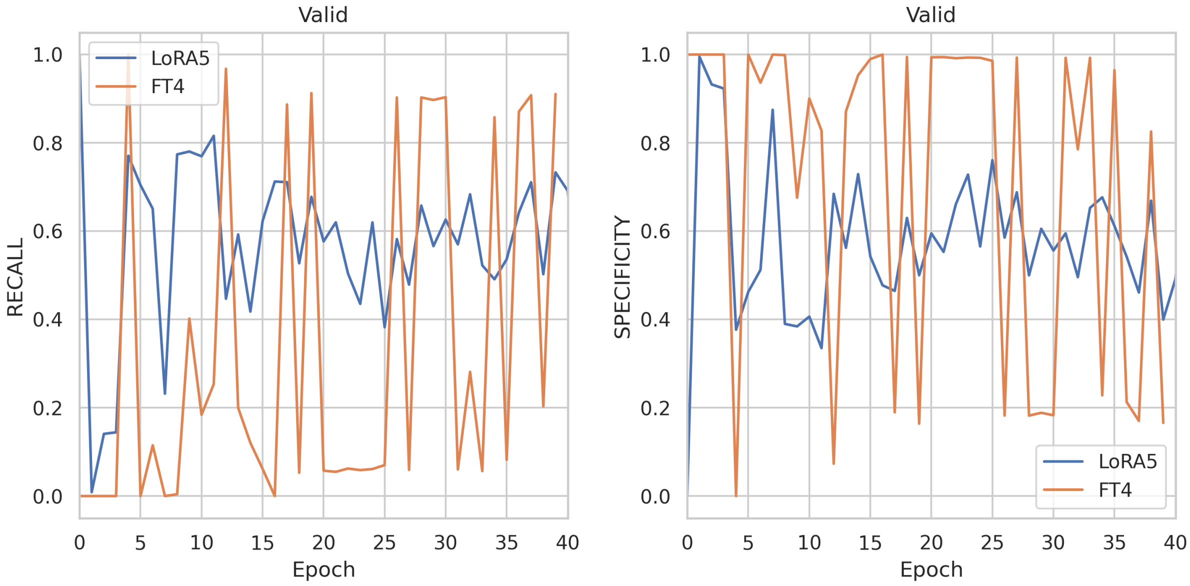


**Fig. S2.** Recall and specificity fluctuate wildly throughout training with traditional fine-tuning, compared to the PEFT model training, which is much more stable. This explains why test set recall is so high, but specificity so low in Table 2—these are highly dependent on the specific epoch which was chosen based on validation AUPR.

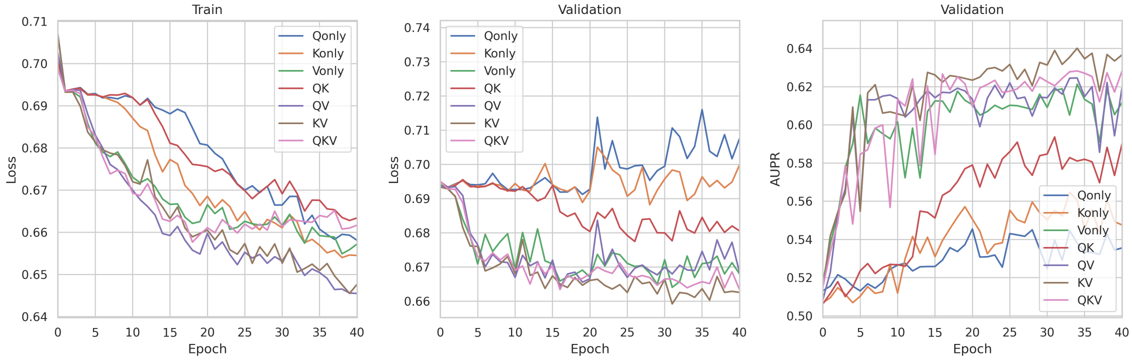


**Fig. S3.** Training and validation loss curves, AUPR curves for models trained with LoRA adapters on $Q$, $K$, $V$, $QK$, $QV$, $KV$, $QKV$ matrices from Table 4. Note that all other training parameters were held constant between these runs.

##### S1.3 Baseline MLP Model

As a baseline to compare with fine-tuning, we train an MLPClassifier model from scikit-learn using embeddings extracted from ESM2 (650M parameters). Parameters for the MLPClassifier were selected by cross-validation on macro average precision over a grid search. We searched over all combinations of

- $activation = [\text{“}logistic''\text{, “}relu''\text{, “}identity'']$
- $alpha = [0.0001, 0.001, 0.01]$
- $learning\_rate\_init = [0.001, 0.01]$
- $max\_iter = 1000, 2000$
- $hidden\_layer\_sizes = [(64, ), (128, ), (512, ), (64, 64), (128, 128), (64, 64, 64)]$
- $tol = [1e - 4, 1e - 5]$

In Figure S5, we show that this MLP model is well calibrated (fraction of predicted positives at threshold $p$ roughly matches predicted probability $p$).

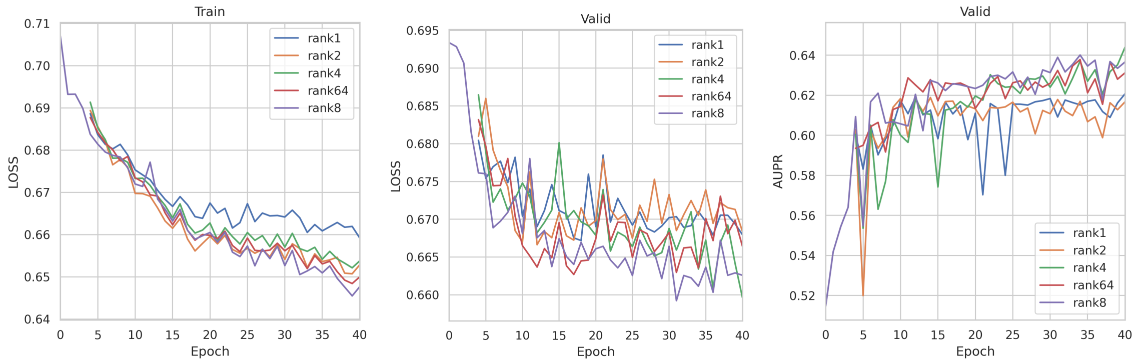


**Fig. S4.** Training and validation loss curves, AUPR curves for models trained with LoRA ranks $r = 1, 2, 4, 8, 64$ from Table 4. Note that all other training parameters were held constant between these runs.

#### S2 Homooligomer Symmetry Prediction

##### S2.1 Test Set Support

Homooligomer symmetry prediction is a highly unbalanced, multi-class prediction task. In Table S2, we show the number of examples in the test set for each class. We show in Figure 2 that PEFT and MLP models are competitive for high support classes like C1, C2, C5, D2, and I, while the FT model is substantially better on rare classes like C7-C9, D4, D5, and O.

**Table S2. Test Set Support.** Number of examples of each symmetry class in the test set. This data is highly imbalanced, with most examples having either C1, C2, D2, or D3 symmetry.

| **Symmetry Class** | **Support** |
|---|---:|
| C1 | 28,899 |
| C2 | 20,671 |
| C3 | 4,666 |
| C4 | 3,057 |
| C5 | 5,955 |
| C6 | 2,885 |
| C7-C9 | 1,406 |
| C10-C17 | 1,910 |
| D2 | 9,384 |
| D3 | 7,539 |
| D4 | 1,377 |
| D5 | 1,700 |
| D6-D12 | 1,975 |
| H | 3,954 |
| O | 520 |
| T | 1,895 |
| I | 4,857 |
| Other | 328 |

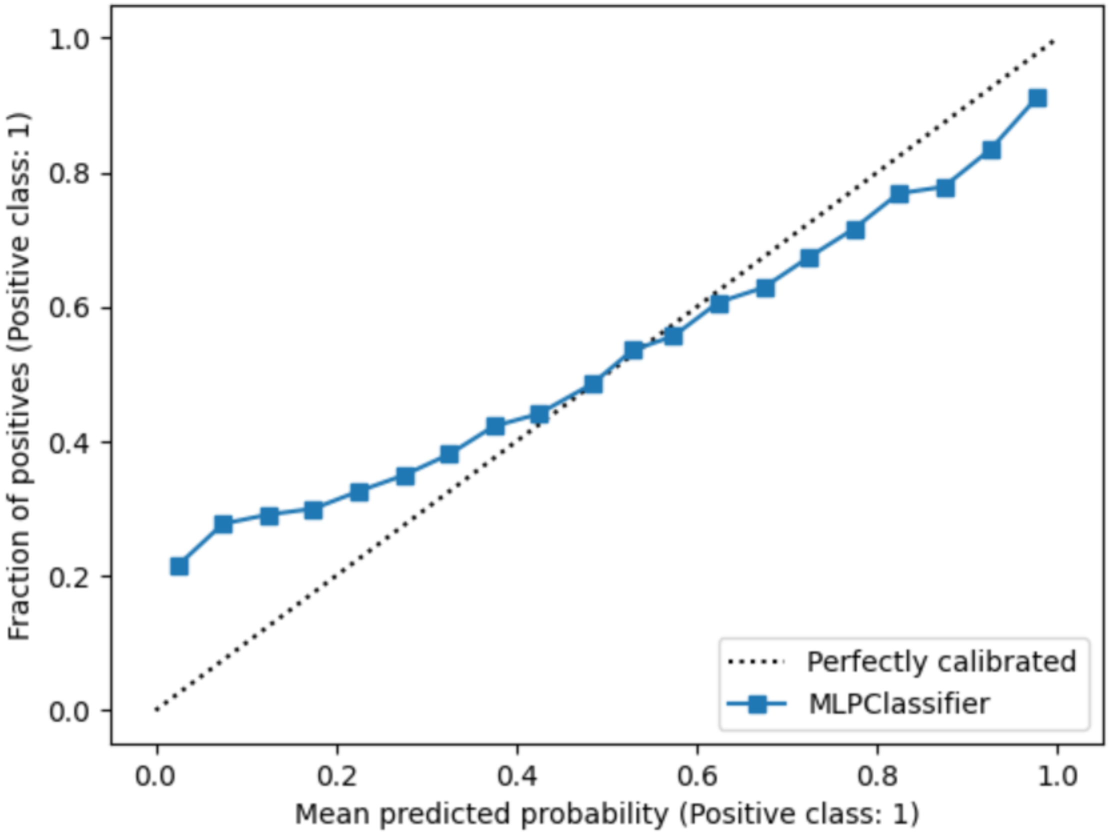


**Fig. S5.** Calibration curve for sklearn MLP on embeddings from a frozen ESM2 model, trained for PPI prediction on the benchmarks from Bernett et al. [8].

##### S2.2 Symmetry Rank Experiments

We show in Table S3 the result of training models with different LoRA rank values for homooligomer symmetry prediction. Here, $r = 8$ is the best performing, closely followed by $r = 4$. Performance drops off noticeably with $r < 4$.

**Table S3. Robustness in rank also holds for homooligomer symmetry prediction.** We perform the same hyper-parameter search as in Section 3.4, this time training models to predict homooligomer symmetry. We find that like for PPI prediction, while performance is respectable at all values, it drops off noticeably for $r < 4$. For symmetry, rank $r = 8$ is the best performing, and there is actually a slight drop off with rank $r = 64$.

| Rank | Val. AUPR | AUPR | Acc. | F1 | MCC | Prec. | Rec. | Spec. |
|---|---|---|---|---|---|---|---|---|
| 1 | 0.506 | 0.359 | 0.345 | 0.352 | 0.428 | 0.419 | 0.345 | 0.968 |
| 2 | 0.525 | 0.385 | 0.351 | 0.369 | 0.450 | 0.516 | 0.351 | 0.969 |
| 4 | **0.531** | 0.403 | 0.372 | 0.383 | 0.455 | 0.503 | 0.372 | 0.969 |
| 8 | 0.461 | **0.416** | **0.390** | **0.430** | **0.468** | **0.558** | **0.390** | **0.970** |
| 64 | 0.445 | 0.388 | 0.359 | 0.358 | 0.420 | 0.444 | 0.359 | 0.968 |

#### S3 Visualizing Attention

We visualize attention values with and without parameter-efficient adapter updates for the last five transformer layers of the PEFT model trained on PPI prediction from Table 2. We average the output of all 20 attention heads so that for a protein of length $n$, we get a matrix of size $n \times n$. LoRA weights are turned on or off with the `peft` package commands `disable_adapter_layers` and `enable_adapter_layers`. We find that with fine-tuning on PPI prediction, attentions are spread out much further from the diagonal, indicating more distal attention necessary for predicting protein-protein interactions. We show representative examples in Figure 3 and Figure S6, a pair of interaction proteins from the NADH dehydrogenase 1 $\beta$ subcomplex. However, this effect is less noticeable in PEFT models trained to predict homooligomer symmetry, where there is a slight diffusion of attention but it still remains largely concentrated along the main diagonal (Figure S7).

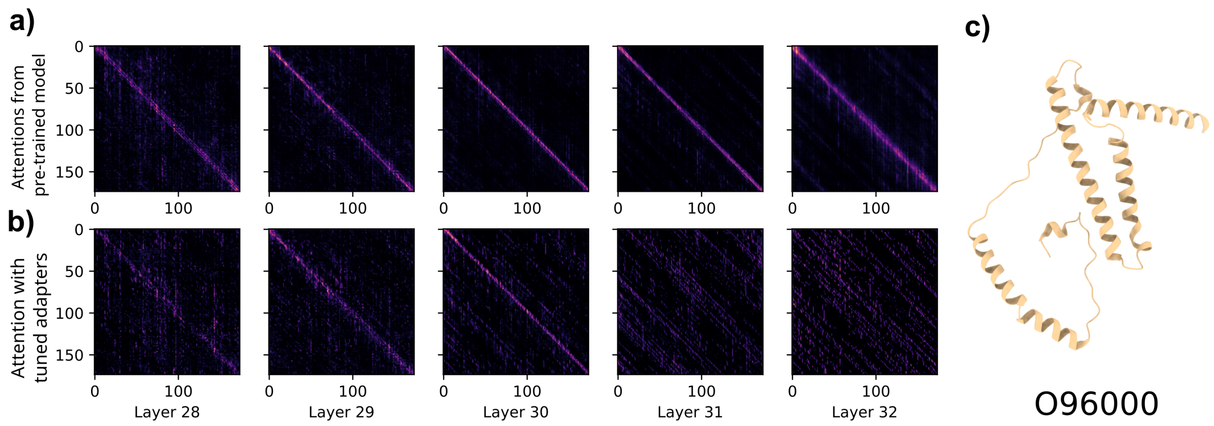


**Fig. S6. Visualizing attention matrices.** **(a)** Attentions for NADH dehydrogenase 1 $\beta$ subcomplex subunit 10 (UniProt ID: O96000) using the pre-trained ESM2. **(b)** Attentions for the same protein after parameter-efficient fine-tuning. **(c)** Structure of O96000. We find that PEFT weights result in attention which is more spread out across the length of the protein when trained for PPI prediction.

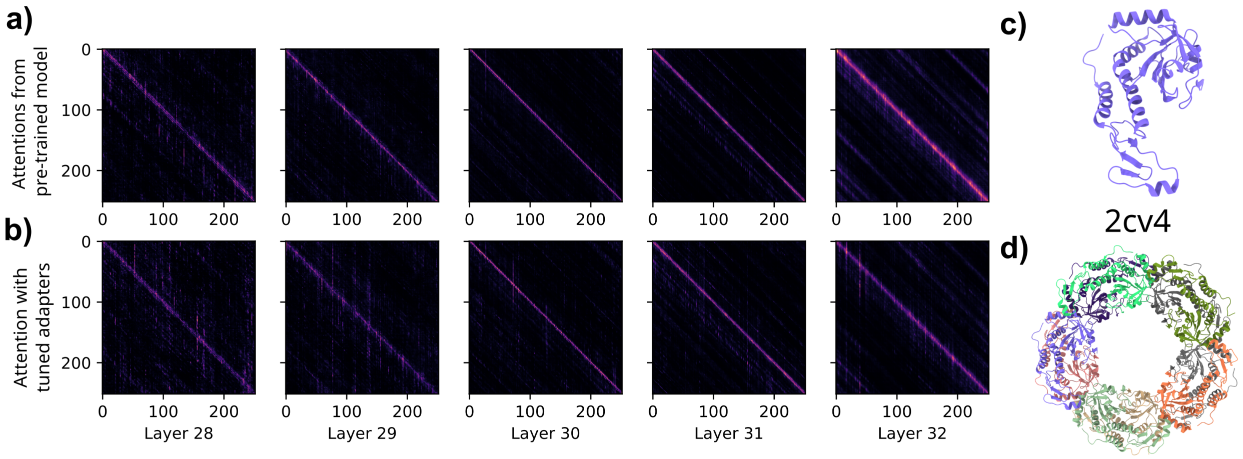


**Fig. S7. Visualizing attention matrices.** **(a)** Attentions for an Archael Peroxiredoxin from the Aerobic Hyperthermophilic Crenarchaeon Aeropyrum pernix K1 (PDB ID: 2CV4), which adopts a dihedral D5 symmetry, using the pre-trained ESM2. **(b)** Attentions for the same protein after parameter-efficient fine-tuning. **(c)** Structure of a single subunit 2CV4. **(d)** Structure of the 2CV4 homooligomer, with D5 symmetry. Here, find that PEFT weights result in attention which is only slight spread more spread across the length of the protein, but still remains concentrated near the diagonal.
